# Incipient speciation, rapid expansion and repeated introgressive hybridization in an African Giant Shrew (*Crocidura olivieri*)

**DOI:** 10.64898/2026.09.02.748917

**Authors:** Malahat Dianat, Violaine Nicolas, Inessa Voet, Josef Bryja, Terrence C. Demos, Christiane Denys, David Ortiz, Julian C. Kerbis Peterhans, Gregory Thom, Adam Konečný, Ondřej Mikula

## Abstract

The African giant shrew (*Crocidura olivieri*) is one of the most widespread native small mammals in sub-Saharan Africa, occupying habitats ranging from humid tropical forests to arid Savanna. Its broad distribution, ecological diversity, and unresolved evolutionary relationships suggest a history of rapid diversification, yet the roles of hybridization and demographic expansion remain poorly understood. We combined genome-wide ddRAD and mitochondrial data from populations sampled across sub-Saharan Africa to reconstruct the evolutionary history of African giant shrews. We reveal extensive mitonuclear discordance across the clade and show that it reflects a complex history of repeated introgressive hybridization rather than a single evolutionary event. Genomic analyses resolve three major evolutionary lineages and identify four introgression events, including extensive bidirectional genome-wide introgression between the ancestors of the *C. olivieri* and arid adapted species followed by later unidirectional introgression associated with range expansion. Within the *C. olivieri* group, we identify five geographically structured lineages connected by ongoing gene flow, representing different stages of incipient speciation. Demographic analyses further reveal repeated population expansion during the Late Pleistocene and Early Holocene, with lineage-specific timing consistent with climatic fluctuations and a possible contribution of increasing human association. Our results identify repeated range expansion and hybridization as major drivers of diversification in African giant shrews and establish this system as a powerful model for studying the genomic processes underlying early speciation.

## Introduction

Species that remain within the same distribution range over long periods of time may become well adapted to the challenges and opportunities of the local environment, especially if it remains stable (1, 2). The opposite evolutionary dynamics is expected when a species expands beyond its previous range, especially if the expansion is rapid and extensive. As larger spatial scales are involved, a greater diversity of abiotic conditions and biotic interactions is encountered. Although such expansion requires a degree of ecological versatility, it also exposes populations to novel selective pressures (3) that drive divergent evolution and potentially promote adaptive radiation (4–7). As a result, new adaptations may arise in different parts of the expanding range; the species may become more specialized at its periphery and may hybridize with newly encountered closely related species (8–10).

*Crocidura olivieri*, one of the most widespread African shrews, appears as a prime example of species undergoing this eco-evolutionary process. In sub-Saharan Africa, *C. olivieri* is one of the most widely distributed native small mammals, occurring from the southern margins of the Congo Basin and montane regions south of Lake Malawi to the Ethiopian Rift Valley, the coastal regions of Senegal, and arid areas of Mauritania. It occupies a broad range of habitats, spanning the forest–savanna continuum from humid closed-canopy forests to seasonally dry savannas and Sudano–Sahelian systems – an unusual ecological breadth for shrews, which are typically associated with moist habitats (11, 12). Its diet is primarily insectivorous, with occasional carrion consumption, but it has also been documented preying on small rodents such as *Mus musculus*, indicating notable trophic flexibility (11, 13). In accordance with these carnivorous habits, *C. olivieri* is the largest African shrew, with body mass ranging from 37 to 65 g (11). In some parts of its range, *C. olivieri* occurs in association with humans (14–16). Its extensive distribution has long been linked to the adoption of a synanthropic lifestyle (11, 17), which may facilitate persistence in human-modified or fragmented landscapes (18–22).

*Crocidura olivieri* belongs to a monophyletic group of large-bodied African shrews often referred to as the “African giant shrews” (23, 24). Previous studies identified two major clades there: *C. olivieri* group, which is widely distributed across sub-Saharan Africa and includes *C. olivieri*, *C. goliath*, *C. viaria*, *C. fulvastra* and *C. somalica* (23, 25, 26); and *C. hirta–flavescens* group, occurring mainly in savanna-like habitats of southern and eastern Africa and including *C. hirta*, *C. flavescens* and *C. bloyeti* (24). The most comprehensive phylogeographic analysis of *C. olivieri* group revealed a complex picture with unresolved nuclear phylogeny, a lot of mismatches between mitochondrial relations and traditional taxonomy and very similar mitochondrial sequences found across large geographic distances (26). This suggests that *C. olivieri* and its close relatives represent an example of early-stage evolutionary radiation, characterized by closely related lineages occupying a wide range of ecological conditions and promoted by rapid geographic spread (11, 26). Such systems are increasingly recognized as valuable models for investigating adaptation and speciation, because rapid geographic expansion exposes lineages to novel ecological conditions (27, 28) and may promote hybridization and introgression at expansion front (e.g. 8, 29).

This study integrates reduced-representation nuclear genomic and mitochondrial data to resolve the phylogenetic structure and divergence history of the African giant shrews across sub-Saharan Africa. We examine how range expansion, secondary contact, and introgression – facilitated by climatic fluctuations and recent human-mediated habitat modification – have shaped lineage diversification and early stages of adaptive radiation.

## Results

### Mitonuclear discordance in African giant shrews

The individual-level nuclear phylogeny of African giant shrews was estimated from single nucleotide polymorphisms (SNPs) concatenated across 1,342 double digest restriction-site associated DNA (ddRAD) loci using Bayesian inference implemented in MrBayes v3.2.7a (30). Basal relationships within the clade were fully resolved, with posterior probabilities (PP) ≥ 0.97 (Figure 1). The phylogeny recovered *C. somalica* as sister to the remainder of the clade, encompassing also individuals originally classified as *C. viaria* and *C. fulvastra*. It therefore encompasses all relatively small-bodied individuals from dry habitats of the Sudano–Sahelian savanna belt. It redefines *C. olivieri* group, therefore, which is now restricted to a pair of large-bodied species, *C. olivieri* and *C. goliath*. This redefined *C. olivieri* group is in sister relationship to *C. hirta*–*flavescens* group, which newly includes also an additional unnamed lineage from Kapiti Plains in Kenya. For a detailed phylogenetic analysis of the whole clade we defined seven species, therefore, which we call *C. somalica*, *C. hirta*, *C. flavescens*, *C. bloyeti*, *C.* sp. “Kenya”, *C. goliath* and *C. olivieri* (Figure 2A).

**Figure 1.**
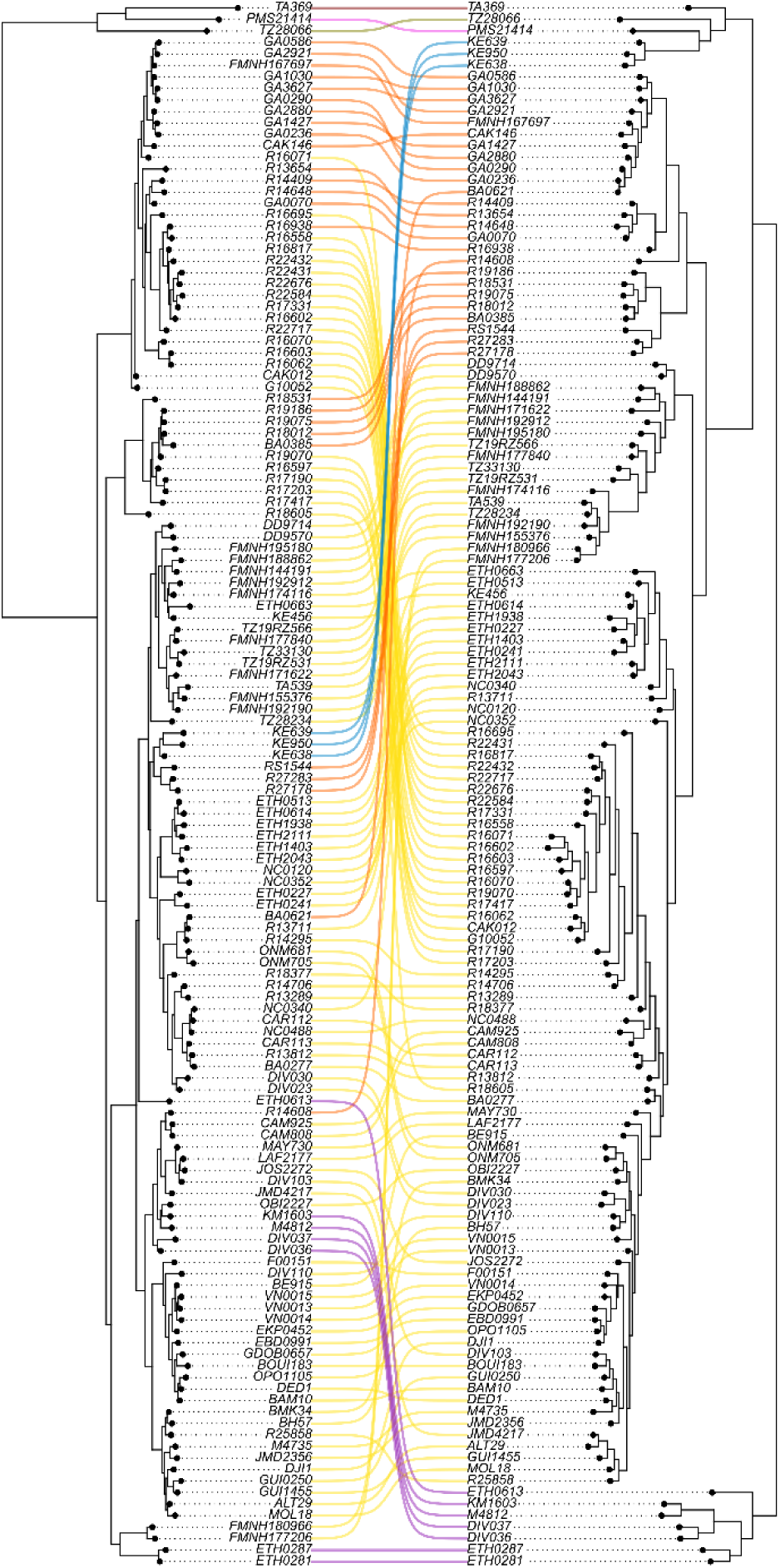
Mitonuclear discordance in African giant shrews. Tanglegram comparing the mitochondrial (left) and nuclear (right) phylogenetic trees. Lines connect the same individuals across the two trees, revealing patterns of mitonuclear discordance. Colours indicate the seven species and correspond to those used in Fig. 2A

**Figure 2.**
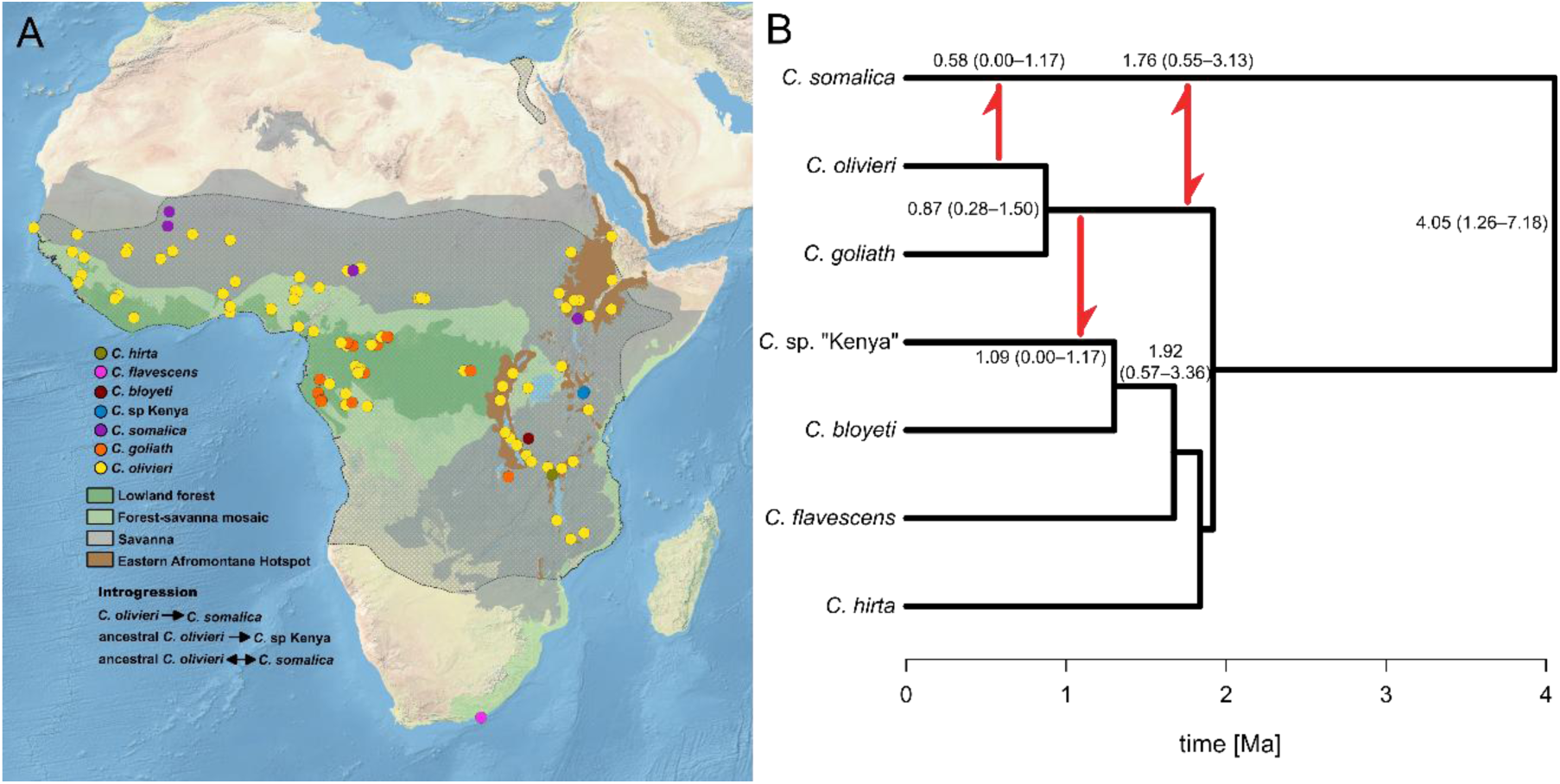
(A) Geographic distribution of the seven *Crocidura* taxa included in this study across major African biotic zones. The grey dotted outline represents the IUCN distribution range of *C. olivieri* from the IUCN Red List (87). Major African biotic zones are based on the terrestrial ecoregions of Dinerstein et al. (88). Arrows indicate the direction of mitochondrial introgression. (B) Time-calibrated phylogeny of seven species of African giant shrews. Red arrows indicate timing and direction of nuclear introgressions. Posterior means and 95% HPD intervals of selected divergence times and introgression events are associated with the respective nodes and arrows.

The mitochondrial (mt) phylogeny was inferred from cytochrome *b* (*CYTB*) sequences in the same way as the nuclear phylogeny. All eight lineages (I-VIII) distinguished by (26) could be found in the *CYTB* tree (Figure 1), along with an additional lineage represented by a single individual (ETH0613). Included is also one representative from each *C. hirta*, *C. flavescens* and *C. bloyeti*. When the two phylogenies were faced against each other, the trees revealed widespread mitonuclear discordance (Figure 1). First, seven individuals of *C. somalica* harbor mtDNA from three different mt lineages, in accord with their geographic origin (Figure S1). Two of them are from south-western Ethiopia (lineage VII), one from North-Western Ethiopia (ETH0613) and four others from Sahel (from lineage III). None of them may be, however, the original mtDNA of the species as none of them is in sister relationship to the rest of the clade and not even to the rest of *C. olivieri* group. Also, all of them have relatively close non-*somalica* relatives in mt tree (<0.05 substitutions / bp). Another case of discordance involves sp. “Kenya”, a member of *C. hirta*-*flavescens* group. Three individuals from the same locality in Kapiti plains were studied and they all carry similar mtDNA, phylogenetically placed deep inside *C. olivieri* group (lineage IV). Finally, there is a discordance within the *C. hirta*-*flavescens* group: *C. bloyeti* is sister to the rest of the group in the nuclear phylogeny but in mt tree it forms sister pair with *C. flavescens*. Extensive discordance was apparent also between *C. goliath* and *C. olivieri*, i.e., over the basal split in the *C. olivieri* group. While in the nuclear tree *C. goliath* is present as a well-defined monophyletic unit (PP=1.00), it contains mtDNA from four different lineages (IV, VI, III, V) (Figure S1) and there was no mitochondrial lineage that would be present exclusively in *C. goliath*.

### Reticulated evolution of African giant shrews

The phylogeny of the seven species was inferred from the ddRAD data using Bayesian multispecies coalescent (MSC) methods implemented in BPP v4.7.0 (31). Four independent runs of the initial analysis, with species boundaries fixed but tree topology estimated, did not converge on the same solution. *Crocidura somalica* was recovered as sister either to the remainder of the clade or to the *C. olivieri* group, despite decisive support (PP=1.00) for both relationships in individual runs. This result suggested insufficient Markov Chain Monte Carlo (MCMC) mixing across topologies; therefore, both alternative topologies were included in subsequent model selection based on marginal likelihoods. Estimates of other relationships in the clade were consistent across the runs, although support for the basal branching order within the *C. hirta*-*flavescens* group was always weak (PP=0.56-0.72).

Marginal likelihoods were estimated in BPP (32) for a series of models, including the pure MSC and the MSC with introgression (MSC-I) (33), with introgression links suggested by instances of mitonuclear discordance. Namely, we considered introgression between *C. olivieri* or ancestor of *C. olivieri* group (“ancestral *C. olivieri*”) on one side and *C. somalica* or *C.* sp. “Kenya” on the other. The fifth possible introgression was between *C. olivieri* and *C. goliath* and all five could lead in both directions, giving the total of ten possible links. We did not assume any ghost lineage to be involved and hence just one time parameter per link was estimated. Initially, two series of models were evaluated using log Bayes factors (BF), i.e., differences in log marginal likelihoods. The series differed in the species tree topology and contained models that included one of ten possible links or no introgression (pure MSC). This comparison provided decisive support for a model with *C. somalica* being sister to the rest of the clade and an introgression involved. In particular, the best supported model included introgression from the ancestral *C. olivieri* into the lineage of *C. somalica*. This model had log BF of 19.19 and 54.95, when compared with, respectively, the best-supported model with the alternative topology and the pure MSC model with the same topology. Both values greatly exceed the threshold of 4.6, which indicates strong evidence (34). Successive inclusion of introgression events led to the final model with four introgression links: introgression from ancestral *C. olivieri* into *C. somalica* was accompanied by introgression in the opposite direction, followed by introgression from ancestral *C. olivieri* into *C.* sp. “Kenya” and from *C. olivieri* to *C. somalica* (Figure 2B). This model received strong to very strong support relative to the best-supported models with one, two, or three introgression links, with log BFs of 12.6, 5.4, and 3.3, respectively.

In the MSC-I model, each introgression event is associated with a pair of introgressions probabilities (φ and 1 − φ), which can be interpreted as the proportions of the genome contributed by the two parental species. In the bidirectional gene exchange between ancestral *C. olivieri* and *C. somalica*, gene flow was balanced: the posterior mean of φ was approximately 0.31 in both directions. In the two later introgression events, from ancestral *C. olivieri* into *C*. sp. “Kenya” and from *C. olivieri* into *C. somalica*, the introgressed proportion was much lower (φ ≈ 0.10; see Table 1 for the estimates). Although the credibility intervals of these estimates are relatively wide, they show little overlap between the bidirectional and the later unidirectional introgression events (Table 1). We therefore conclude that the ancient hybridization event was substantially more extensive.

**Table 1.** Estimates of introgression times (rescaled to Ma) and probabilities (φ) in the best-supported MSC-I model. For both parameters, posterior mean and 95% HPD interval limits are given.

| Donor species | Acceptor species | Time (Ma) | Introgression probability ($\phi$) |
| --- | --- | --- | --- |
| *C. somalica* | ancestral *C. olivieri* | 1.76 (0.55 – 3.13) | 0.316 (0.143 – 0.470) |
| ancestral *C. olivieri* | *C. somalica* | 1.76 (0.55 – 3.13) | 0.315 (0.162 – 0.481) |
| ancestral *C. olivieri* | *C. sp. "Kenya"* | 1.09 (0.31 – 1.93) | 0.122 (0.053 – 0.206) |
| *C. olivieri* | *C. somalica* | 0.58 (0.00 – 1.17) | 0.097 (0.019 – 0.177) |

For time calibration of the resulting reticulated tree, we performed a divergence dating analysis in StarBEAST 2 (35) using an *ad hoc* assembled ddRAD sequence dataset of species from across phylogeny of the genus and calibration prior, which approximated the root age of *Crocidura* from (36). The divergence of *C. olivieri*–*C. goliath* was dated to 0.87 (0.28 – 1.50) Ma and it was then used to scale all times in the MSC-I species tree (Figure 2B). In particular, the basal split between *C. somalica* and the remainder of the clade was dated to 4.05 (1.25 – 7.18) Ma while the introgression events took place from 1.76 to 0.58 Ma (Table 1).

### Population structure of *C. olivieri* group

The coancestry matrix of *C. olivieri* group indicates moderate population structure without large off-diagonal blocks of uniformly low coancestry indicating long-isolated lineages (cf. 37). The fineSTRUCTURE analysis identified 41 clusters (Table S1, Figure S2) in a dendrogram where five major geographically distinct groups were distinguished (Figure 3A, Table S1). We therefore interpret these five groups as phylogeographic lineages and designate them Old Central, New Central, Western, Northeastern and Eastern (Figure 3B). The Western and Northeastern lineages span the entire longitudinal extent of the Sahelo-Sudanian savanna, extending eastward to the Ethiopian sector of the Great Rift Valley (GRV). The Old Central lineage is so named because it occurs in the Congo Basin and corresponds to *C. goliath*; that is, it is sister to the remainder of the group. In contrast, the New Central lineage, which also occurs in the western Congo Basin, is phylogenetically nested within the remaining *C. olivieri* lineages. Finally, the Eastern lineage is largely restricted to montane regions along the equatorial sector of GRV. The distribution ranges partly overlap; especially the Old Central lineage is at some localities found together with New Central, Northeastern and Eastern lineages (Figure 3B).

**Figure 3.**
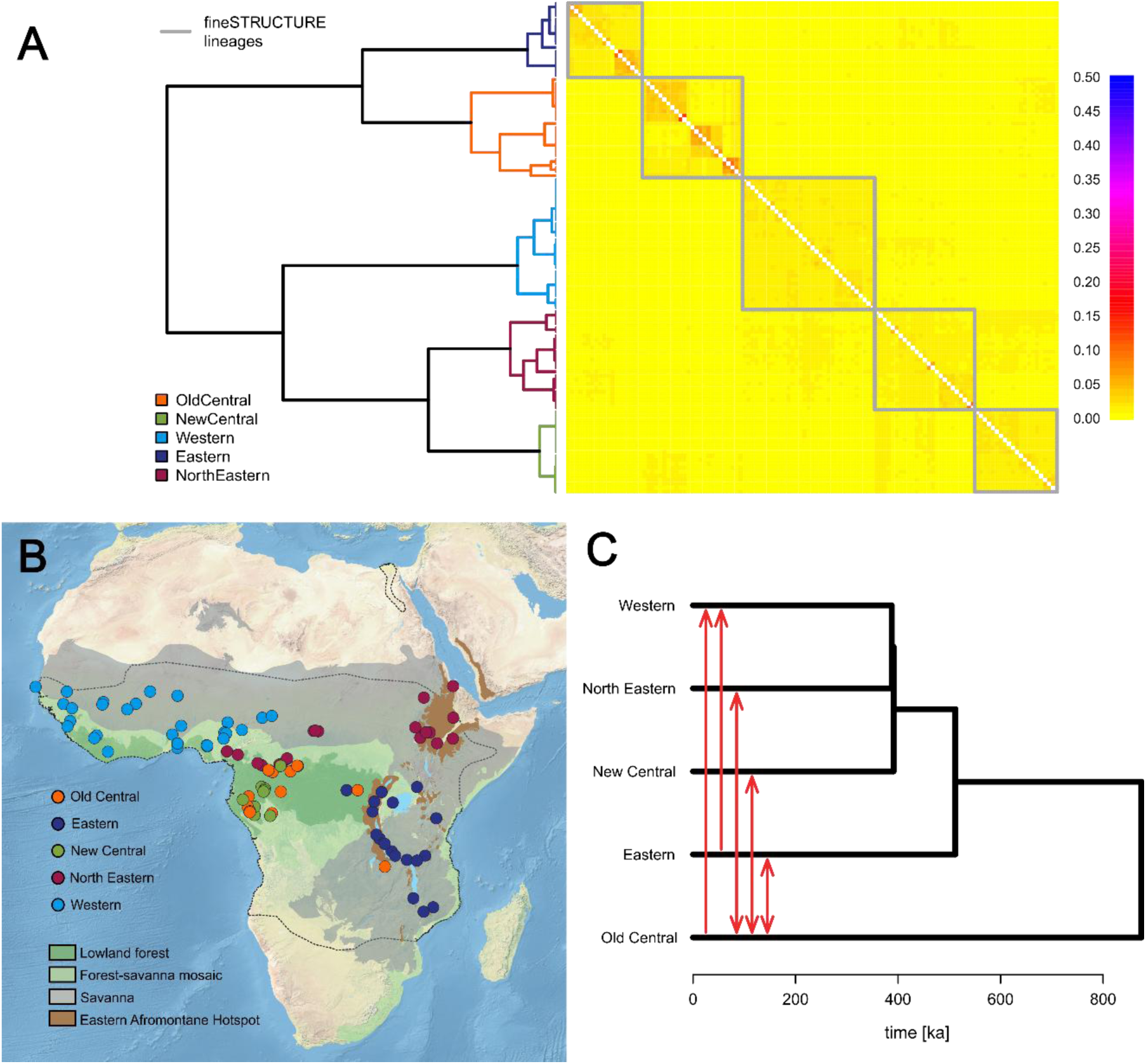
(A) Co-ancestry matrix and corresponding dendrogram showing relationships among the inferred populations and their assignment to the five major nuclear lineages. (B) Geographic distribution of the five fineSTRUCTURE lineages (Old Central, Eastern, New Central, Northeastern, and Western) across major African biotic zones (based on the terrestrial ecoregions of Dinerstein et al. (88)). The grey dotted outline represents the IUCN distribution range of *C. olivieri* (87). (C) Time-calibrated MSC-M tree of the five nuclear lineages. Red arrows indicate the direction of gene flow. Arrow positions do not indicate the timing of gene flow; gene flow applies to the period during which both branches coexisted. The time axis is given in thousands of years (ka).

In the space defined by the first two principal components, the lineages formed coherent clusters, that were tight, elongated towards the origin of the axes or showing some subdivision (Figure S3). However, some individuals from Old Central, Eastern and Western were substantially shifted towards the origin, where they may overlap. In the unrooted tree estimated in MrBayes from concatenated ddRAD data (one SNP per locus), the five lineages were also almost completely separated – only the westernmost samples of the Northeastern were closer to either New Central or Western samples (Figure S3).

Differentiation of the lineages was assessed in a pairwise manner. First, admixture analyses in STRUCTURE were run with K = 2 and no prior information on cluster membership. These analyses supported the lineages as at least partially distinct gene pools, as they consistently estimated a large proportion of individual genotypes to be derived purely from one of the ancestral populations and high interclass heterozygosity (H_IC_) was associated with large divergence of ancestral allele frequencies (D). At the same time, although H_IC_ varied considerably, a proportion of admixed individuals was always present (Table 2).

**Table 2.** Estimates of inter-lineage admixture based on pairwise analyses. The ancestral frequency divergence (D) and inter-class heterozygosity (H_IC_) from STRUCTURE is supplemented by migration rates (M) or divergence time (*τ*) from BPP. If non-zero values of MSC-M parameters were not supported (Savage-Dickey ratio ≪ 100), the respective cell is left empty. The direction of migration is indicated by an arrow symbol.

| Lineages | D | $H_{IC}$ | M→ | M← | $\tau$ |
| --- | --- | --- | --- | --- | --- |
| Old Central ↔ Eastern | 0.0931 | 0.0228 | 0.0422 | 0.0857 | 0.0022 |
| Old Central ↔ New Central | 0.0518 | 0.0037 | 0.0839 | 0.1657 | 0.0016 |
| Old Central ↔ Western | 0.0634 | 0.0097 | 0.0320 | 0.1057 | 0.0022 |
| Old Central ↔ Northeastern | 0.0866 | 0.0117 | 0.0501 | 0.1123 | 0.0020 |
| Eastern ↔ New Central | 0.0974 | 0.0282 | 0.0342 | 0.0414 | 0.0018 |
| Eastern ↔ Western | 0.1279 | 0.0345 | 0.0443 | 0.0470 | 0.0017 |
| Eastern ↔ Northeastern | 0.1291 | 0.0590 |  |  | 0.0014 |
| New Central ↔ Western | 0.0895 | 0.0135 | 0.0511 | 0.0524 | 0.0014 |
| New Central ↔ Northeastern | 0.1385 | 0.0953 |  |  | 0.0011 |
| Northeastern ↔ Western | 0.1208 | 0.0317 |  |  |  |

Next, MSC with migration (MSC-M) model implemented in BPP (38) was used to co-estimate divergence time (*τ*) and the rate of continuous migration between lineages (M). Significance was tested using the Savage-Dickey (SD) density ratio (39). Almost all lineage pairs were found to be separated by significantly non-zero divergence times, and in most cases non-zero bidirectional migration was detected (Table 2). SD ratios distinguished zero and non-zero parameters very neatly as they were either ≤ 3 (no support) or > 100 (usually ≫100), indicating very strong support.

The two analyses thus provided complementary perspectives and, at first glance, seemingly contradictory results regarding admixture. The oldest split – between Old Central and the remaining lineages – was accompanied by the highest estimates of gene flow (particularly into, rather than from, Old Central) and, paradoxically, relatively low estimates of D and H_IC_. In contrast, where divergence was shallow to zero and M was non-significant, D and H_IC_ were among the highest. When integrated, these results show that even relatively old lineages are capable of interbreeding, but even the most admixed populations differ in their ancestral allelic frequencies. In the BPP analyses, each lineage was represented by a random sample of alleles form across its distribution, which allowed post-divergence gene flow between older lineages to be distinguished from shared ancestral polymorphism. In STRUCTURE analyses, alleles were not a priori assigned to any lineage and the use of all individual genotypes accounted for geographical variation. This allowed to detect large differences in allelic frequencies, presumably driven by geographically distant samples, together with substantial admixture where the lineages were in contact. On the other hand, relatively intensive gene flow between predefined lineages could diminish population-level divergence and individual-level admixture when they are co-estimated without a priori set lineage boundaries. Thus, between more divergent and geographically less widespread lineages, e.g., two ‘Central’ lineages, relatively few individuals were found to be of highly admixed ancestry.

Overall phylogeographic picture of *C. olivieri* group was obtained by applying STRUCTURE and BPP analyses to all five lineages together. The degree of admixture revealed by STRUCTURE with K = 5 was much higher compared to the pairwise estimates (SI). The overall H_IC_ was 0.0931, almost as high as the maximum pairwise estimate (between New Central and Northeastern lineages) and substantially higher than the others. Accordingly, the individual-level admixture showed much broader transitions between the lineages than in most pairwise analyses.

The final phylogeny of *C. olivieri* group was inferred in BPP under MSC with migration (MSC-M) model with eight significant post-divergence gene flows detected by means of Savage-Dickey (SD) density ratio (39) in five pairs of lineages (Figure 3C). In three of them, the migration was bidirectional, namely between the Old Central lineage and three others (New Central, Northeastern and Eastern). The others were unidirectional, from Western to Old Central and from Eastern to Western lineages. The phylogeny fully parametrized MSC-M species tree showed that divergences between the lineages were generally very shallow relative to population size. Only the length of Old Central branch approached unity in coalescent units, all others were substantially shorter. In particular, the divergence of New Central, Northeastern and Western lineages is very young and virtually polytomic. Extensive sharing of genes is therefore expected in the *C. olivieri* group, both due to migration and incomplete lineage sorting (ILS).

### Population trends in *C. olivieri* group

Our demographic analyses rejected the neutral model for all populations. The two-epoch model (2Pins), indicating substantial historical population growth, provided the best fit (lowest AIC) for all populations except the Northeastern population, which was best explained by the three-epoch model (4Pins), suggesting an initial moderate increase followed by rapid expansion (Table S2; Figure S4). Using estimated θ values, we converted parameters to absolute units to infer population sizes and timing of changes. Overall, the Northeastern population exhibited the greatest expansion. All populations experienced demographic expansion. The Eastern population expanded to 7.3-fold around 15 ka, the New Central population 11.9-fold around 23 ka, the Old Central population 4.7-fold around 15 ka, and the Western population 12.2-fold around 7 ka. In contrast, the Northeastern population underwent two successive expansion phases, with an initial 3.2-fold increase around 33 ka followed by a second expansion around 16 ka, resulting in an overall 30-fold increase in population size (Table S3).

## Discussion

### Diversity of African giant shrews

The evolutionary relationship between *Crocidura olivieri* and morphologically similar but smaller-bodied, arid-adapted forms remained unresolved in the previous studies based on a limited number of mitochondrial and nuclear markers (26). Our genomic analyses reveal a complex story of reticulated evolution that gave rise to three major evolutionary lineages. The basal divergence is about twice as old as the following ones and occurred between the ancestors of the *C. somalica* group and all remaining species. The next one happened between the *C. hirta–flavescens* and *C. olivieri* groups, approximately in the Pliocene or early Pleistocene, and it was soon followed by extensive introgression between the newly diverged *C. olivieri* lineage and the *C. somalica* lineage. If not synanthropic, the *C. olivieri* group inhabits forests or forest-savanna mosaic (26). Both its earliest branches (Old Central and Eastern) occur in rainforests of the Congo Basin or in the moist habitats of the eastern African highlands which suggests the group expanded from the Central African lowland forests into a wide variety of habitats across sub-Saharan Africa. At the same time, the ancestral *C. olivieri* was not probably restricted to distinctly humid environments as it was involved in introgressive hybridization with the other two species groups whose species inhabit drier and more open habitats (24, 26) which therefore likely represent their ancestral condition.

The inference based on genome-scale data and discovery of widespread mitonuclear discordance revises the species diversity of African giant shrews. It supports a single species in the *C. somalica* group, but at least four in the *C. hirta–flavescens* group. Both these groups inhabit similar kind of habitats and marginally overlap in southern Ethiopia. While *C. somalica* inhabits relatively uniform Sudanian savanna belt, however, the *C. hirta–flavescens* group lives in landscapes of the eastern and southern Africa, that are topographically complex due to presence of the East African Rift, large lakes, and the Eastern Afromontane highlands. This could increase diversity of suitable habitats on the large geographic scale and also make their climate-driven fragmentation more likely, both factors increasing probability of speciation.

### The history of introgressions

The most striking feature of the evolutionary history of African giant shrews are the recurrent interspecific introgressions. The inferred events ranged from extensive bidirectional nuclear exchange between ancestral lineages to later unidirectional nuclear introgressions and geographically localized mitochondrial capture. The most extensive introgression occurred when *C. somalica* and the ancestral *C. olivieri* came into contact, most probably during the early Pleistocene (∼1.8 Ma before present). The gene flow was bidirectional and, in each direction, it involved a substantial fraction of genome (ca. 31% of genome). Later (∼0.6 Ma), the already introgressed genome of *C. somalica* received another contribution (ca. 10%) from the modern *C. olivieri*. Apart from that, we documented three independent instances of mitochondrial introgression from *C. olivieri* to *C. somalica* as the introgressed haplotypes occur at three different positions in the mitochondrial phylogeny. In West Africa, the introgression was apparently recent, but some of the two Ethiopian mitochondrial transfers might correspond to the younger nuclear introgression. None of the introgressed haplotypes form a deep-branching lineage that could be attributed to the ancient bidirectional exchange. Finally, there is both nuclear and mitochondrial introgression into *C.* sp. “Kenya”, but, strikingly, these two were likely distinct events. The nuclear introgression was from the ancestral *C. olivieri*, while the introgressed haplotypes are nested among other *C. olivieri* haplotypes and the mitochondrial capture therefore had to occur more recently.

The reasons and consequences of introgression can also vary across the events. The ancient bidirectional introgression might happen simply due to the intensity of contacts between broadly distributed and – in that time – relatively recently diverged species. It is also possible, however, that the introgressed material increased the adaptive potential of the gene pools and facilitated the expansion of the *C. olivieri* group across the forest–savanna gradient (e.g. 40, 41). Distinguishing these alternatives will require whole-genome sequence data to identify introgressed regions and test whether they show evidence of selection or contain genes plausibly linked to ecological adaptation. Generally, range expansion provides a plausible setting for further hybridization, which can also explain repeated mitochondrial capture. It does not explain, however, why mtDNA consistently moved from *C. olivieri* into dry-adapted taxa.

Given that mitochondrial protein-coding genes are crucial for energetic metabolism, accepting foreign mtDNA can be adaptive, if it provides a better functional match to conditions near species’ range margins (42–44). In that case, however, we would expect introgression in the opposite direction as it is *C. olivieri* for which the increasingly dry environment on its distribution margins should be challenging. Expansion models also generally predict resident mtDNA introgressing into an expanding lineage particularly under male-biased dispersal (45, 46). This expectation could be reversed, however, if the resident dry-adapted taxa occurred in small, fragmented populations and the expanding *C. olivieri* outnumbered them locally. Small populations may also accumulate mildly deleterious mitochondrial mutations (47, 48), which can favor their replacement by heterospecific haplotypes with a lower mutational load (49). Female-biased dispersal – reported in *C. russula* (50) – could reinforce introgression into the resident species, because expanding females of *C. olivieri* would carry its mtDNA into them.

### Drivers of population expansions in *C. olivieri* group

Remarkably, every major lineage within the *C. olivieri* group bears the signature of demographic expansion. The inferred onsets of lineage expansions (ca. 33–7 ka) must be interpreted with some caution due to uncertainty in divergence dating, but together they form a chronological sequence spanning two contrasting climatic phases – from the Last Glacial Maximum (LGM; ca. 23–19 ka) to the African Humid Period (AHP; ca. 15.2–5.5 ka). The LGM is generally associated with rainforest fragmentation and grassland expansion (51), but its effects were spatially heterogeneous across the central Africa: forests contracted in some regions but persisted or expanded in others, often as open forest–woodland mosaics (52, 53). Subsequent postglacial humidification promoted vegetation recovery, culminating in the AHP with the continent-wide intensification of monsoon precipitation (54, 55).

*Crocidura* shrews are generally more abundant in moist habitats (11) and hence unlikely to expand when the climate is getting drier. On the other hand, the African Giant shrews as a whole and the *C. olivieri* group in particular show remarkable ecological versatility, which can provide them with competitive advantage in conditions that are suboptimal for most other shrews. The inferred expansions can be, therefore, associated either with a local expansion of forest-savanna mosaic or recovery of humid conditions. The expansion of the New Central lineage at the onset of LGM (ca. 23 ka) might be an example of the former, while expansions of the Northeastern, Eastern and Old Central lineages at the onset of the AHP (ca. 15-16 ka) might exemplify the latter. The most recent of the expansions, that of the Western lineage dated to ca. 7 ka, could be also reinforced by increasing association with humans, whose population expanded 3–10 ka in western Africa (56).

### Synanthropy and persistence in arid regions

The Sahelo-Sudanian savanna belt represents gradual transition between the Guineo-Congolian forests and the Sahara Desert. Given the general association of shrews with relatively moist habitats, *C. olivieri* occurs surprisingly far in the dry parts of the gradient. Especially the Western lineage extends into arid regions of Mali, Senegal, Burkina Faso, and Niger, while additional *C. olivieri* specimens from Mauritania indicate that the species reaches even farther into the Saharan margin (Z. Boratyński, personal communication). Available distributional records strongly suggest this is possible due to adoption of commensal lifestyle with human settlements providing buffered microhabitats and additional food resources. In southern Senegal, *C. olivieri* was frequently captured inside buildings (16), consistent with earlier records from across the western Africa (26). Available observations from the arid northern Senegal are also predominantly from human dwellings rather than from natural habitats (A. Dalecky, personal communication). In Nigeria, *C. olivieri* was abundant in settlements and cultivated habitats but rare in natural rainforest and savanna, suggesting that human-modified habitats promoted its occurrence (17).

Similar patterns were observed in other small mammals as well. The murine rodent *Praomys daltoni* occurs mainly in human settlements in the arid Sahelian habitats, which also suggests that synanthropy compensates for otherwise unsuitable climatic conditions (57, 58). Also, the tropical Asian house shrew *Suncus murinus* is largely restricted to human-associated habitats around coastal settlements in the arid Arabian Peninsula (59).

### Phylogeographic structure of *C. olivieri* group

In any case, the *C. olivieri* group is strongly geographically structured. In this study, we adopted a five-lineage model based on a fineSTRUCTURE dendrogram, which reflects shared patterns of individual coancestry and thus can identify appropriate basic units for population genetic analyses. It is not a phylogenetic estimate but under conditions of recent or ongoing diversification, populations may not be structured into a strict hierarchy of fully discrete units.

The lineage formation can be understood from results of pairwise admixture analyses (Table 2). Although admixture coefficients of STRUCTURE estimate contributions of ancestral gene pools, not literally a degree of post-divergence admixture, a straightforward expectation is that inter-class heterozygosity from STRUCTURE will be positively correlated with migration rates from BPP. We observed the opposite, however. The highest Ms (e.g., New Central → Old Central lineage) are associated with low H_IC_ values. The converse is also true as the non-significant (virtually zero) Ms are observed where H_IC_ is the highest (e.g., New Central ↔ Northeastern).

This may reflect two different modes of divergence. First, some lineage pairs appear to have experienced a period of isolation followed by secondary contact. The intervening isolation produced a clearer divergence signal and a larger estimated tau, allowing BPP to distinguish post-divergence migration from ILS. At the same time, when admixture is restricted to narrow contact zones, H_IC_ remains low despite a relatively high estimated M. This pattern was observed between Old Central and each of the New Central, Northeastern, and Western lineages. In contrast, the Northeastern–Western and Northeastern–New-Central pairs may have diverged gradually through isolation by distance as an ancestral population expanded in different directions. Serial founder events during expansion can rapidly shift allele frequencies, producing substantial overall differentiation (high D) without generating a consistent genealogical split. Because these lineages did not share a distinct period of isolation, their estimated divergence times (tau) are very small, so BPP cannot distinguish migration from ILS and estimates M as nonsignificant. However, populations close to the inferred origin of expansion retained extensive ancestral allele sharing, producing high H_IC_.

In terms of the actual geography, the primary divergence of the Western–Northeastern–New-Central triad took place probably in the forest–savanna mosaic along the northern margin of the Congo Basin, encompassing parts of Cameroon, northern Democratic Republic of Congo, and the Central African Republic. It can also explain small differences between fineSTRUCTURE lineages and both the ordination in the first two principal components and the branching pattern of the concatenated tree. The discrepancy is confined to six individuals of the Northeastern lineage from the putative area of primary divergence and can be therefore explained by sharing of ancestral polymorphism and its different impact on different analyses. Finally, it is also good to note that although the Eastern lineage seems to be geographically well separated from the Old Central lineage, there is actually a sampling gap between them. They can be in contact as indicated by our two samples of the Eastern lineage from the Congo basin – and there can be a good deal of gene flow between them.

### Incipient speciation in *C. olivieri* group

Old Central was formally described as *C. goliath* based on its reported larger size (26). Here, its status as a separate species was supported by its degree of genetic differentiation, but the ongoing gene flow suggests that its reproductive isolation from *C. olivieri* is at least incomplete. The MSC-M analyses supported gene flow between the Old Central lineage (*C. goliath*) and all four lineages of *C. olivieri*. Effective migration rates were consistently two-to threefold higher toward *C. goliath* than in the opposite direction. Given that the two most divergent lineages both occur in the Congo Basin and that one of them (Old Central) is internally structured there, we infer that this region represents the ancestral area. This implies that gene flow is again preferentially from expanding populations to the native one, consistent with the lower estimated effective population size of the Old Central lineage. The estimated migration rate was approximately 0.1, or one immigrant per ten generations, an order of magnitude lower than the one-migrant-per-generation threshold expected to counteract neutral genetic differentiation caused by genetic drift (60). Nevertheless, this rate is sufficient for even moderately advantageous alleles to spread across the species boundary (61). Overall, the reproductive barrier between *C. olivieri* and *C. goliath* appears porous but sufficiently strong to keep the two species genetically distinguishable even in syntopy.

The nature of the reproductive barrier remains unknown. However, a degree of ecological segregation is suggested by habitat information available for 19 and 21 individuals from the Old Central (*C. goliath*) and New Central (*C. olivieri*) lineages, respectively. Among them, 14 *C. goliath* individuals were collected in primary forest whereas sympatric-to-syntopic *C. olivieri* occupied a broader range of habitats, including primary and secondary forests, human-modified forests, savannas, and forest patches within savanna landscapes. Even such partial segregation can reduce genetic exchange between populations (62) and contribute to differentiation despite ongoing gene flow (63, 64).

### Implications and Outlook

The evolutionary history of the *C. olivieri* group is remarkable not for any single process, but for their convergence: recurrent introgression, rapid expansion and diversification across the forest–savanna gradient, possible human-assisted expansion into and persistence at the Saharan margin, and emerging species boundaries maintained despite ongoing gene flow. By bringing these processes together within a single continental radiation, African giant shrews provide a powerful natural experiment for understanding how hybridization, climatic change, and human modification interact to generate and maintain biodiversity.

Population-scale whole-genome sequencing could determine whether introgression from dry-adapted congeners supplied adaptive variation during the colonization of arid environments. Paired sampling of natural and anthropogenic habitats across the aridity gradient, combined with physiological and microclimatic measurements, could distinguish intrinsic heat and desiccation tolerance from environmental buffering provided by human settlements. The persistence of genetic differentiation between *C. goliath* and New Central *C. olivieri* despite ongoing gene flow provides a tractable system for studying incipient speciation. Standardized sampling across primary, secondary, and anthropogenic habitats, integrated with morphology, and trophic ecology, could test whether ecological and phenotypic differentiation restricts admixture and maintains their distinctiveness.

## Materials and Methods

### Sequence data

In total, we analyzed sequence data from 140 individuals including samples from across the geographic range of the *C. olivieri* group (Figure 2A) as well as from its close relatives (*C. hirta*, *C. flavescens*, *C. bloyeti* and *C. somalica*) and four outgroup species (*C. yaldeni*, *C. lamottei*, *C. cyanea*, and *C.* cf. *hildegardeae*). The specimens come from the collections of CBGP: Centre de Biologie pour la Gestion des Populations, Baillarguet, Montferrier-sur-Lez, France. FMNH: Field Museum of Natural History, Chicago, USA. IVB: Institute of Vertebrate Biology, Studenec, Czech Republic. IZEA: Institut de Zoologie et d’Ecologie Animale, Lausanne, Switzerland. MNHN: National Museum of Natural History, Paris, France. MVZ: Museum of Vertebrate Zoology, Berkeley, USA. NHM: Natural History Museum, London, UK. PMS: Slovenian Museum of Natural History, Ljubljana, Slovenia. RBINS: Royal Belgian Institute of Natural Sciences, Brussels, Belgium. ZFMK: Zoologisches Forschungsmuseum Alexander Koenig, Bonn, Germany. For the comprehensive list of material, see Table S1.

Reduced representation nuclear genomic data were obtained by double digest restriction site associated DNA (ddRAD) sequencing. The genomic libraries were prepared following the protocol of (65) and the sequencing performed commercially using the Illumina NextSeq 550 Mid-Output Kit (150 bp paired-end reads) at the Genomics Core Facility of CEITEC (Brno, Czech Republic) and the NovaSeq PE150 platform at Novogene (UK). Raw sequence data are available in the NCBI Sequence Read Archive under the accession number XXXXX (Table S1). After demultiplexing and quality filtering of raw reads (detailed in Text S1), alignments of ddRAD loci were obtained in ipyrad v0.9.61 (66). We used its *de novo* assembly method with 85% similarity threshold, the minimum required sequencing depth of 6x and other filtering criteria set as detailed in Text S1. Based on an exploratory analysis of sequencing success (Text S1) we retained only loci with at least 70% occupancy (sequencing success across individuals). Importantly, the assembly was performed and filtered separately with either all 140 individuals (‘full’ data set) and just 123 of them, belonging to the *C. olivieri* group (‘*olivieri’* data set). The ‘full’ data set contained 1,342 loci with 37,764 variable sites (i.e., single nucleotide polymorphisms or SNPs). Out of these, 1,340 loci contained at least one SNP which was biallelic and parsimony informative, which means its minor allele was present in at least two copies. The ‘*olivieri’* data set contained 2,148 loci with 46,467 SNPs in total. The biallelic parsimony informative SNPs were present at 2,142 loci. For some analyses, we used ‘single SNP’ versions of these data sets, which means we retained just a single SNP per locus. For modelling of demographic trends, we just required the SNPs to be biallelic, otherwise they were selected at random. For other analyses we also constrained the selection only to parsimony informative SNPs. Generally, we used sequences phased in two parental haplotypes per individual at each locus (“.alleles” output of ipyrad), although for some analyses we subset or collapsed the haplotypes as explained in descriptions of particular methods.

Finally, we analyzed also an alignment of cytochrome *b* (*CYTB*) gene sequences (402–1140 bp long), which represented mitochondrial DNA of the same 140 ddRAD-sequenced individuals. Of these, 67 sequences (including the outgroups) were downloaded from NCBI GenBank, and 73 sequences were newly generated (following the protocol of (67)) and deposited in GenBank under accession numbers XXXX– XXXX (Table S1).

### Phylogeny of *C. olivieri* and its relatives

The phylogeny of the *C. olivieri* group and its close relatives was estimated, and mitonuclear discordance was inspected using the full data set. First, Bayesian nuclear and mitochondrial trees were inferred from concatenated single SNP and *CYTB* sequences, respectively. Both the ddRAD and *CYTB* trees were inferred as unrooted, with branch lengths expressed in substitution units, and their inference followed the same procedure. Nucleotide substitution models – and, in the case of *CYTB*, a codon-partitioning scheme – were pre-selected using ModelFinder (68) in IQ-TREE v2.2.0 (69). The tree inference was then performed in MrBayes v3.2.7a (30), with posterior density distribution sampled using Markov Chain Monte Carlo (MCMC). Convergence of four independent runs was assessed using the potential scale reduction factor (70) and the average standard deviation of split frequencies (71). Posterior samples were pooled after discarding the initial 10% of each as burn-in. The pooled posterior sample of trees was then used to estimate the maximum clade credibility (MCC) topology (72) and the posterior probabilities of its bipartitions. Branch lengths were then estimated as posterior means from a second MrBayes analysis with identical settings, but with the topology fixed. The resulting MCC tree and posterior sample (from the unconstrained topology analysis) were rooted using the outgroups, which were subsequently pruned. The SNP genotypes are phased within loci but not between them and so the concatenation required collapsing two parental haplotypes into a single sequence with heterozygous positions represented by ambiguity codes.

In the ddRAD tree, seven species were recognized (see Results) and their phylogeny further analyzed in the framework of multispecies coalescent (MSC) model with introgression (MSC-I, (33)) in BPP v4.7.0 (31). Furthermore, each species was represented by two randomly chosen sequences at each locus. In all these analyses, the topology of the backbone species tree was fixed to that of the concatenated MCC tree and weakly informative priors were put on divergence time (*τ*) and population size (*θ*) parameters (see Text S2 for the details). The pure MSC model and a series of introgression (MSC-I) scenarios were tested. The specification of introgression links, each with its associated introgression probability (*φ*) was motivated by patterns of mitonuclear discordance (see Results). The models were compared by their log marginal likelihoods estimated by the path sampling method with 32 points bridging the prior and posterior distributions (32, 73). The ratio of marginal likelihoods – or, equivalently, the exponential of the difference between the log marginal likelihoods of two models – is the Bayes factor (BF), which quantifies the strength of evidence in favor of the better-supported model. Values of log BF ≥ 3 (or BF ≥ 20) are typically interpreted as strong evidence, while log BF ≥ 4.6 (or BF ≥ 100) means very strong evidence (34). Introgression links were first tested individually and, if significant, sequentially added to the model in decreasing order of significance, with each addition evaluated as part of an increasingly complex model.

### Population structure of *C. olivieri* group

Genomic similarities of individuals in the ‘*olivieri*’ data set were first explored by the clustering on the coancestry matrix, estimated from complete haplotype sequences. Each row of this matrix contains counts of loci at which haplotypes of the individual *i* were found most similar to the haplotypes of particular individuals *j* ≠ *i*. The matrix is therefore of dimension *N* × *N* with zeros on its diagonal and each of its elements *C_i_*_,*j*_ approximates frequency of the first order coalescence relationships between individuals *i* and *j* across local gene trees (74). For the clustering, we used fineSTRUCTURE, which partitions individuals to the smallest possible clusters with distinct coancestry profiles, i.e. mean coancestry between members of such clusters (75). The clustering is based on the multinomial distribution model and performed in a Bayesian framework, so its result is represented by the maximum a posteriori state. Then, the hierarchical clustering is performed, which successively joins clusters whose merging causes the smallest drop in log-likelihood. Branch lengths in the resulting dendrogram correspond to these log-likelihood differences.

The population structure of the *C. olivieri-goliath* group was further explored by two analyses of the single SNP data set. First, we used principal component analysis (PCA) as described in (76). Secondly, we inferred a phylogenetic tree from the concatenated alignment exactly as from the ‘full’ data set, but without outgroups, so the tree was left unrooted.

### Pairwise admixture analyses

Further analysis of population structure in *C. olivieri-goliath* group was built upon the fineSTRUCTURE analysis, which allowed to define five distinct lineages (see Results). Their mutual relationships were explored by pairwise analyses based on two complementary population genetic models. The admixture model of STRUCTURE (77) with two ancestral populations (K=2) assumes individual genotypes are made up by contributions of two ancestral populations. These are characterized by different allelic frequencies and genotypes in the corresponding Hardy-Weinberg proportions. Their contributions are quantified by individual admixture coefficients *Q*, which are co-estimated together with the ancestral allelic frequencies. No prior information on cluster membership was used. The population-level admixture (*A_p_*) was quantified as a mean interclass heterozygosity, 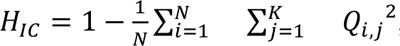, i.e., a probability that two alleles randomly picked from an individual are of different ancestry. The net divergence in ancestral allele frequencies (*D*) was estimated and reported by STRUCTURE as described in (78). The MSC model with migration (MSC-M), as implemented in BPP v4.7.0 (38), explicitly models continuous migration – rather than episodic introgression – occurring between populations since their divergence. The migration rate is quantified in units of the effective number of migrants per generation (*M*) and can be asymmetric – that is, absent or unequal in one or both directions. Its significance was assessed using Savage-Dickey (SD) density ratio, computationally less demanding approximation of BF, which we used to test for non-zero migration rate (39).

### Phylogeographic scenario

The final set of genetic analyses was carried out to provide a basis for outlining the overall phylogeographic scenario. First, an admixture analysis in STRUCTURE was performed with K = 5, implicitly assuming that each lineage primarily descends from a distinct ancestral population. Next, we used BPP to build an increasingly complex MSC-M model by incrementally adding post-divergence migrations connecting branches of the species tree. The procedure was similar as previously applied when adding introgression links. However, we already tested these migrations in pairwise analyses, so they were added into the model in decreasing order of their significance. Also, we added them to the model as bidirectional and only when shown insignificant in one or both directions, they were dropped. Finally, we used SD ratio (instead of path sampling) to test for significance of migrations. All STRUCTURE analyses (pairwise and overall) were conducted on the parsimony informative single SNP data matrix. All BPP analyses were conducted on two sequences per lineage per locus. In these analyses we also used just a subset of 2,127 loci, which were successfully sequenced in all five lineages.

We inferred historical population size changes for each of the five distinct lineages using dadi (79), which models site frequency spectra (SFS) following the dadi-pipeline workflow described by (80). SNP datasets were converted to SFS using dadi.Misc.make_data_dict in dadi. To maximize the number of segregating sites while accommodating missing data, SFS were projected to reduced sample sizes for each population following (80). We applied standard demographic models implemented in dadi (Demographics1D.py), including, the 0P model corresponds to a constant population size, 2Pins and 2Pexp describe instantaneous and exponential size changes, 3P represents a bottleneck followed by growth, and 4Pins and 4Pexp describe two successive size changes, instantaneous and exponential, respectively (81). For model optimization, we performed five rounds with 20, 20, 30, 20, and 5 replicates, and maximum iterations of 1, 5, 20, 50, and none per round, respectively. Grid sizes were set to 58, 68, and 78, and parameter perturbation factors to 3, 2, 1, and 0.5 across rounds. Parameter estimation was conducted using the Nelder–Mead algorithm. For each population, only the replicate with the highest log-likelihood was retained for model comparison, and competing models were ranked using the Akaike Information Criterion (AIC; (82)), with the lowest AIC indicating the best fit (80). To convert the model parameters (*nu* and *t*) into population size (*N*) and time in years (*T*), we used θ estimated by dadi to calculate the ancestral effective population size (N_ref_) (83). N_ref_ was computed as *N_ref_*= *θ*/(4*Lμ*), where *L* is the number of SNPs and *μ* is the per-site mutation rate. We used a mutation rate of 1.41 × 10⁻⁹ per site per generation (assuming a generation time “g” of one year), estimated by comparing branch lengths in substitution units from the MSC-I tree (0.0012) with divergence times from the StarBEAST analysis (0.87 Ma) between *C. olivieri* and *C. goliath* (see Results). Using this mutation rate, the demographic parameters were finally converted according to the expressions *N* = *nu* × *N_ref_* and *T* = 2*t* × *N_ref_* × *g* (84).

The time scale of the phylogeographic events was set by rescaling of the MSC-I and MSC-M species trees and the time axis of population trends. The time calibration information was obtained by divergence dating analysis using an *ad hoc* assembled ddRAD sequence dataset that included 46 individuals from 12 species from across phylogeny of the genus. Included were five species from this study (*C. olivieri*, *C. goliath*, *C. hirta*, *C. flavescens* and *C. somalica*) so the results were informative about divergences and clock rates in *C. olivieri* clade, but the tree was not overly unbalanced. The ddRAD loci were assembled in ipyrad v0.9.61 with the same parameter settings as before, but using mapping to reference, which was a genome of *Crocidura gueldenstaedtii* (GenBank Accession GCA_965212715.2). Raw data for this task were taken from this and previously published studies (24, 85). The assembly resulted in 182 loci that were represented in at least 11 out of 12 species and we further randomly selected 30 of them to meet computational demands of the analysis. The analysis was conducted in MSC framework as implemented in StarBEAST2 (35), a package of BEAST v2.7.5 (86). Using MSC accounts for the difference between divergence times in the species tree and in the gene trees and makes the estimated dates directly applicable to the MSC-I and MSC-M. The estimate of the root age of *Crocidura* was taken from (36) and translated into the calibration prior with gamma distribution (*α* = 2.0, *β* = 0.9 and offset=13.36), which has mean=15.6 and 95% limits 13.6 and 19.6, in units of Ma. We assumed strict molecular clock in the analysis and put a broad uninformative prior on its rate. Tree topology was constrained to that of Dubey et al. (25) and this study. For relative divergence times we assumed homogenous birth-death process with uninformative priors. Population size parameters were analytically integrated out in the inference. In all loci, we assumed HKY nucleotide substitution process with locus-specific rates.

## Acknowledgments

This study was supported by the ATM Blanche 2019 of the MNHN (Paris, France), the project of the Czech Science Foundation no. 23-06116S, and the Czech-French Contact Mobility – Barrande Program (Project PHC Barrande 46665SB and MSMT 8J21FR008). This work was supported from Operational Programme Research, Development and Education – “Project Internal Grant Agency of Masaryk University” (No. CZ.02.2.69/0.0/0.0/19_073/0016943).

Acknowledgments go out to numerous individuals who contributed to the fieldwork, including R. Šumbera, J. Šklíba, J. Goüy de Bellocq, V. Mazoch, T. Aghová, Y. Meheretu, D. Mizerovská, M. Uhrová, O. Mikula, L. Cuypers, T. Zewdneh, J. Zima jr, A. Ribas, M. Lövy, S. Šafarčíková, J. Krásová, J. Kreisinger, K. Welegerima, A. Didier Missoup, P. Barrière, M. Colyn, Akora, Ch. Sabuni and all local collaborators. Our gratitude goes to Z. Boratynski for providing us with a sample of *Crocidura olivieri* from Mauritania, marking the first documented record of this species in the country. We would like to show our appreciation to J.C. Kerbis Peterhans, T. Demos and A. Ferguson from Field Museum of Natural History, Chicago, USA (FMNH), L. Granjon from Centre de Biologie pour la Gestion des Populations, Montferrier-sur-Lez, France (CBGP), E. Verheyen from Royal Belgian Institute of Natural Sciences, Brussels, Belgium (RBINS), A. Olayemi from Natural History Museum, Obafemi Awolowo University, Ile Ife, Nigeria, S. Gambalemoke Mbalitini from University of Kisangani, Faculty of Sciences, Laboratory of Ecology and Management of Animal Resources (LEGERA), Kisangani, DR Congo, J. Patton from Museum of Vertebrate Zoology, Berkeley, USA (MVZ), B. Kryštufek from Slovenian Museum of Natural History, Ljubljana, Slovenia (PMS) for providing us with museum samples.

Computational resources were provided by the e-INFRA CZ project (ID:90254), supported by the Ministry of Education, Youth and Sports of the Czech Republic.

We would like to thank O. Mikula and M. Mahmoudi for their assistance in conducting bioinformatic analysis. We would like to express our thanks to A. Bryjová and D. Čížková for their assistance in preparing the genomic library.

