## supplementary information for "Incipient speciation, rapid expansion and repeated introgressive hybridization in an African Giant Shrew (*Crocidura olivieri*)"

**Malahat Dianat**[1,2]**, Violaine Nicolas[3], Inessa Voet[3], Josef Bryja[1,2], Terrence C. Demos[5,6] Christiane Denys[3], David Ortiz[4], Julian C. Kerbis Peterhans[5,6], Gregory Thom[7], Adam Konečný[1]*, and Ondřej Mikula[2]*
[1] *Department of Botany and Zoology, Faculty of Science, Masaryk University, Brno, Czech Republic*
[2] *Institute of Vertebrate Biology of the Czech Academy of Sciences, Brno, Czech Republic*
[3] *Institut de Systématique, Évolution, Biodiversité (ISYEB), Muséum national d'Histoire naturelle, CNRS, Sorbonne Université, EPHE, Université des Antilles, CP 51, 75005 Paris, France*
[4]*Museum of Zoology, University of Puerto Rico at Río Piedras, San Juan, Puerto Rico*
[5] *Negaunee Integrative Research Center, Field Museum of Natural History, Chicago, Illinois, USA*
[6] *College of Arts and Sciences, Roosevelt University, Chicago, Illinois, USA*
[7] *Museum of Natural Science and Department of Biological Sciences, Louisiana State University, Baton Rouge, LA, USA*

* These authors share senior authorship
** Corresponding author: Malahat Dianat

**Figures**

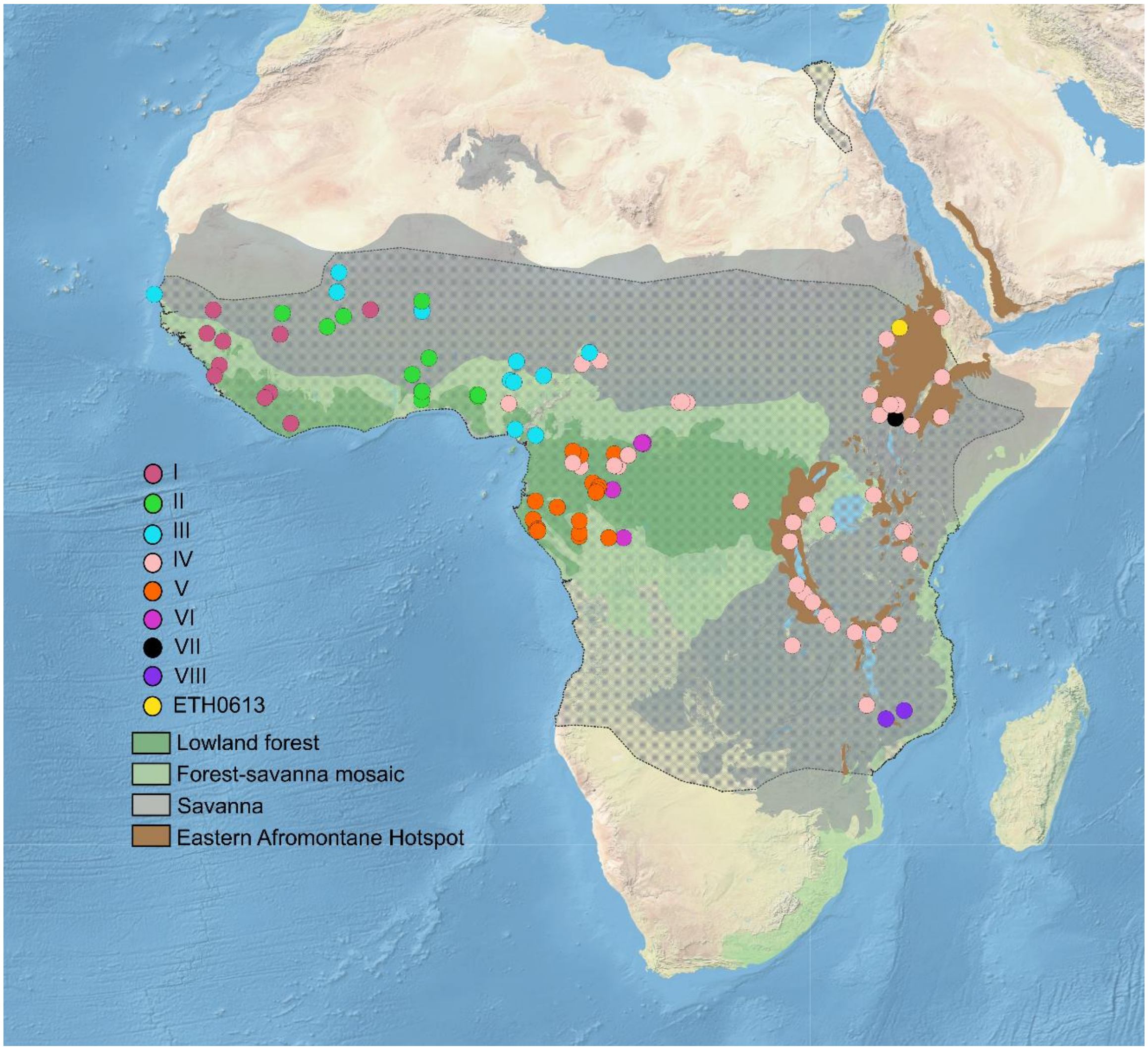


**Figure S1**. Geographic distribution of the nine mitochondrial lineages of Crocidura olivieri across major African biotic zones. The grey dotted outline represents the distribution range of C. olivieri from the IUCN Red List (1). Major biotic zones are based on the terrestrial ecoregions of Dinerstein et al. (2).

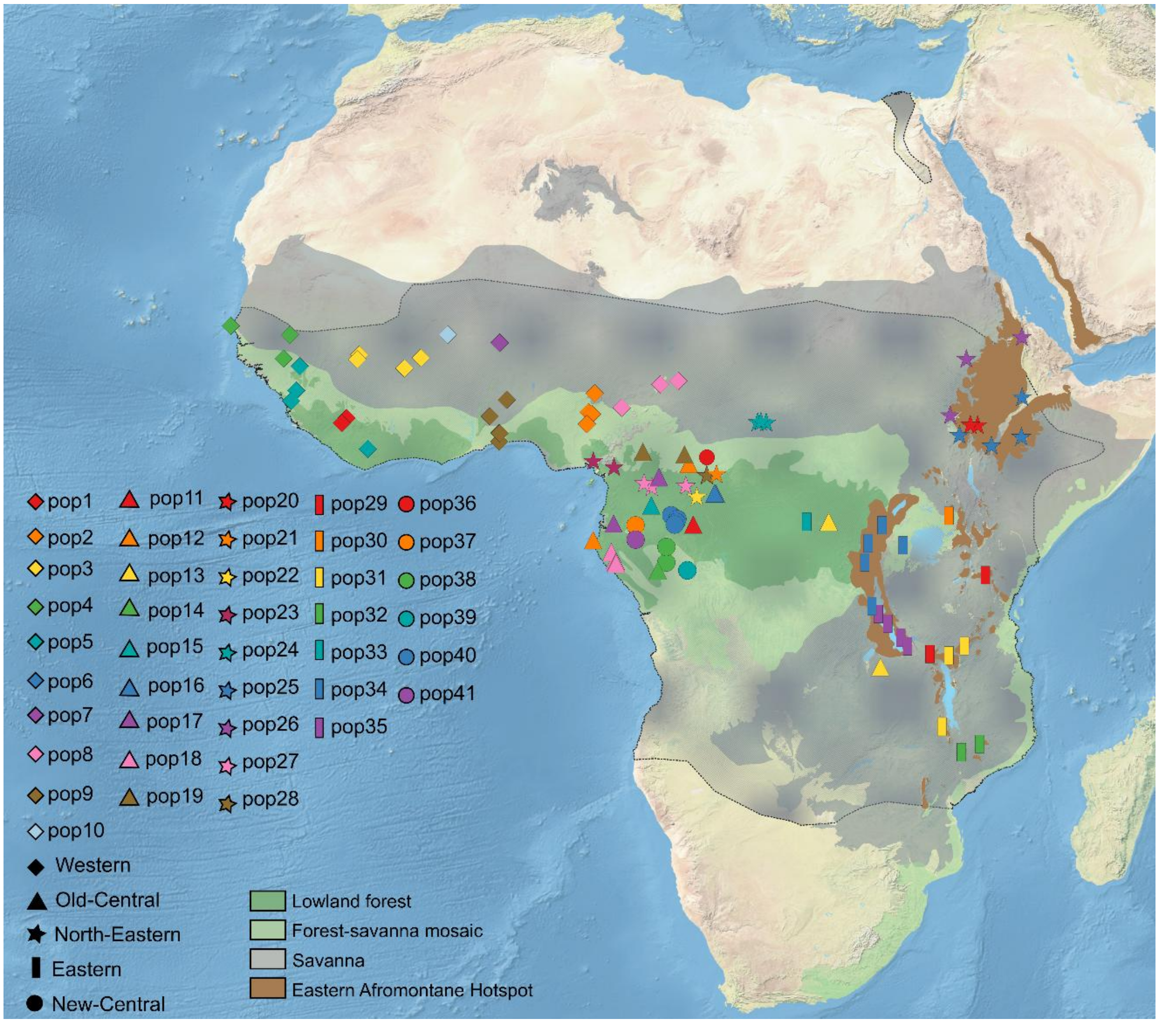


**Figure S2**. Geographic distribution of the 41 population clusters identified by fineSTRUCTURE within the C. olivieri group. Colours and symbols indicate the 41 fineSTRUCTURE clusters (pop 1 – pop41), the symbol shapes indicate to which of the five lineages the clusters belong. The grey dotted outline represents the distribution range of C. olivieri from the IUCN Red List (1). Major African biotic zones are shown in the background and are based on the terrestrial ecoregions of Dinerstein et al (2).

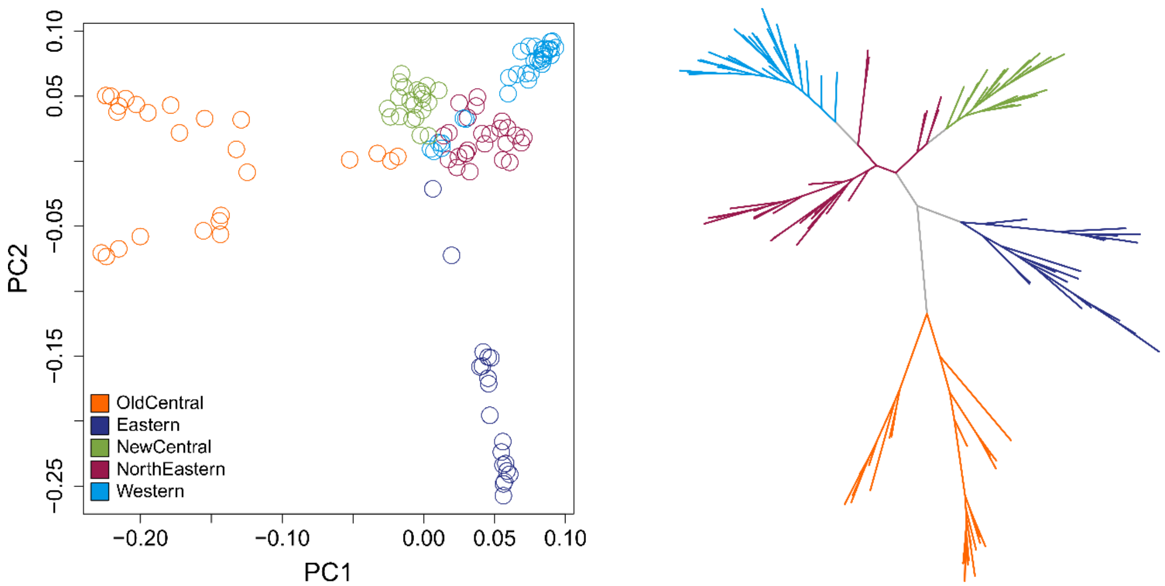


**Figure S3.** Genetic differentiation among the five phylogeographic lineages of the *Crocidura olivieri* group based on ddRAD data. (A) Principal component analysis (PCA) showing the distribution of individuals along the first two principal components. (B) Unrooted Bayesian tree estimated in MrBayes from concatenated ddRAD data using one SNP per locus.

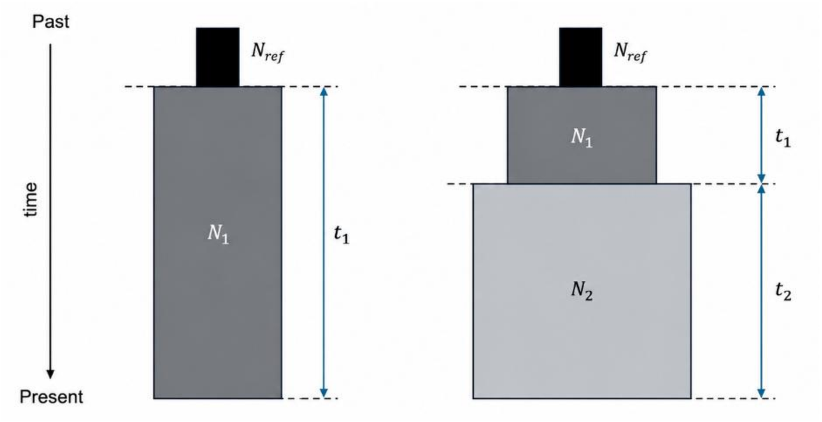


**Figure S4**. Schematic representation of demographic models fitted in dadi for the five populations of the *Crocidura olivieri* group. The two-epoch model (2Pins; single instantaneous size change) on the left, provided the best fit for the Eastern, New-Central, Old-Central, and Western populations, whereas the three-epoch model (4Pins; two instantaneous size changes) on the right, was selected for the North-Eastern population. Population sizes are parameterized as N1 and N2 relative to the ancestral size (Nref), and the timing of changes is given by t1 and t2.

**Tables**

**Table S1.** (Provided as a separate Excel file) Data on *Crocidura* specimens used in this study. CBGP: Centre de Biologie pour la Gestion des Populations, Baillarguet, France. FMNH: Field Museum of Natural History, Chicago, USA. IVB: Institute of Vertebrate Biology, Studenec, Czech Republic. IZEA: Institut de Zoologie et d'Ecologie Animale, Lausanne, Switzerland. MNHN: National Museum of Natural History, Paris, France. MVZ: Museum of Vertebrate Zoology, Berkeley, USA. NHM: Natural History Museum, London, UK. PMS: Slovenian Museum of Natural History, Ljubljana, Slovenia. RBINS: Royal Belgian Institute of Natural Sciences, Brussels, Belgium. ZFMK: Zoologisches Forschungsmuseum Alexander Koenig, Bonn, Germany. Morphometric measurements are included.

**Table S2**. Comparison of demographic models inferred using dadi for each population within the *Crocidura olivieri* group. Akaike Information Criterion (AIC) values are shown for all models, with the best-supported model for each population (lowest AIC) indicated in bold.

| | Eastern | New_Central | North_Eastern | Old_Central | Western |
|---|---|---|---|---|---|
| 2Pins | **73.96** | **65.62** | 75.16 | **85.58** | **84.8** |
| 2Pexp | 75.62 | 67.18 | 75.7 | 86.4 | 86.5 |
| 4Pexp | 75.98 | 69.4 | 75.12 | 89.24 | 88.6 |
| 4Pins | 76.76 | 69.4 | **74.7** | 89.24 | 88.6 |
| 3P | 77 | 67.6 | 75.54 | 87.22 | 86.74 |
| 0P | 180.12 | 247.68 | 405.94 | 160.9 | 284.76 |

**Table S3.** Demographic parameter estimates inferred with dadi for five populations of the *Crocidura olivieri* group. θ is population-scaled mutation parameter. Parameters nu1 and nu2 represent population sizes relative to the ancestral population size, whereas t1 and t2 represent the durations of demographic epochs in coalescent units. Absolute effective population sizes and epoch durations in years were estimated using the mutation rate (μ) and generation time (g). Parameters nu2 and t2 apply only to the North-Eastern population under the three-epoch model, in which the timing of the older demographic event corresponds to the cumulative duration of T1 + T2.

| | Eastern | New_Central | Old_Central | North_Eastern | Western |
|---|---|---|---|---|---|
| number of individuals | 19 | 25 | 25 | 19 | 35 |
| L (number of SNPs) | 468231 | 468231 | 468231 | 468231 | 468231 |
| θ | 43.52 | 25.35 | 47 | 30.52 | 26.05 |
| nu1 | 7.2572 | 11.8845 | 4.6931 | 3.2404 | 12.1521 |
| nu2 | – | – | – | 30 | – |
| t1 | 0.449 | 1.2121 | 0.4138 | 0.707 | 0.6982 |
| t2 | – | – | – | 0.7097 | – |
| μ (per site mutation rate) | 1.41E-09 | 1.41E-09 | 1.41E-09 | 1.41E-09 | 1.41E-09 |
| $N_{ref}= \theta/4L\mu$ | 16445 | 9579 | 17760 | 11533 | 9844 |
| $N1 = nu1 \times N_{ref}$ | 119346 | 113844 | 83351 | 37371 | 119622 |
| $N2 = nu2 \times N_{ref}$ | – | – | – | 345985 | – |
| $T1 = 2 \times t1 \times N_{ref} \times g$ | 14768 | 23222 | 14698 | 16307 | 13746 |
| $T2 = 2 \times t2 \times N_{ref} \times g$ | – | – | – | 16370 | – |
